# Genomic and transcriptomic determinants of therapy resistance and immune landscape evolution during anti-EGFR treatment in colorectal cancer

**DOI:** 10.1101/448076

**Authors:** Andrew Woolston, Khurum Khan, Georgia Spain, Louise J Barber, Beatrice Griffiths, Reyes Gonzalez Exposito, Yatish Patil, Sonia Mansukhani, Matthew N Davies, Andrew Furness, Francesco Sclafani, Clare Peckitt, Kyriakos Kouvelakis, Romana Ranftl, Ruwaida Begum, Isma Rana, Janet Thomas, Annette Bryant, Sergio Quezada, Andrew Wotherspoon, Nasir Khan, Nikolaos Fotiadis, Teresa Marafioti, Thomas Powles, Fernando Calvo, Sebastian Guettler, Katharina von Loga, Sheela Rao, David Watkins, Naureen Starling, Ian Chau, Anguraj Sadanandam, David Cunningham, Marco Gerlinger

**Affiliations:** Translational Oncogenomics Lab, The Institute of Cancer Research, London; GI Cancer Unit, The Royal Marsden Hospital, London; Systems and Precision Cancer Medicine Lab, The Institute of Cancer Research, London; UCL Cancer Institute, University College London; Tumour Microenvironment Lab, The Institute of Cancer Research, London; Department of Radiology, The Royal Marsden Hospital; Departments of Pathology and Histopathology, University College Hospital, London; Barts Cancer Institute, Queen Mary University, London; Division of Structural Biology, The Institute of Cancer Research, London

## Abstract

Anti-epidermal growth factor receptor (EGFR) antibodies (anti-EGFR-Ab) are effective in a subgroup of patients with metastatic colorectal cancer (CRC). We applied genomic and transcriptomic analyses to biopsies from 35 *RAS* wild-type CRCs treated with the anti-EGFR-Ab cetuximab in a prospective trial to interrogate the molecular resistance landscape. This validated transcriptomic CRC-subtypes as predictors of cetuximab benefit; identified novel associations of *NF1*-inactivation and non-canonical *RAS/RAF*-aberrations with primary progression; and of *FGF10*- and non-canonical *BRAF*-aberrations with AR. No genetic resistance drivers were detected in 64% of AR biopsies. The majority of these had switched from the cetuximab-sensitive CMS2-subtype pretreatment to the fibroblast- and growth factor-rich CMS4-subtype at progression. Fibroblast supernatant conferred cetuximab resistance *in vitro*, together supporting subtype-switching as a novel mechanism of AR. Cytotoxic immune infiltrates and immune-checkpoint expression increased following cetuximab responses, potentially providing opportunities to treat CRCs with molecularly heterogeneous AR with immunotherapy.

## MAIN TEXT

Anti-epidermal growth factor receptor (EGFR) antibodies (anti-EGFR-Ab) are effective in a subgroup of patients with metastatic colorectal cancer (CRC). Activating *KRAS* or *NRAS* mutations in codons 12, 13, 59, 61, 117 and 146 have been associated with primary resistance in randomized trials and anti-EGFR-Ab-treatment should only be administered for tumors that are wild-type (wt) at these loci^1-5^. In spite of this stratification, many patients do not benefit, indicating additional resistance mechanisms. *BRAF* V600E^6^, *MAP2K1/ERK1^7^* or *PIK3CA*^8^ mutations, amplifications of *KRAS*^9^ and of the receptor tyrosine kinases (RTKs) *ERBB2*, *MET* and *FGFR1*^7^ have been suggested as further drivers of primary resistance. Moreover, a recently defined transcriptomic classification of CRCs into distinct subtypes found an association of the transit amplifying (TA)-subtype with cetuximab sensitivity^10^, suggesting that non-genetic molecular characteristics also influence anti-EGFR-Ab sensitivity.

Acquired anti-EGFR-Ab resistance almost invariably occurs in patients who initially benefit and this has predominantly been studied retrospectively in circulating tumor DNA (ctDNA) using targeted genetic analyses^11-13^. *KRAS* and *NRAS* mutations, as well as *EGFR*-exodomain mutations that alter the binding epitope for the anti-EGFR-Ab cetuximab have been found in a large proportion of patients with acquired resistance. Amplifications of *MET* or *KRAS* evolved in some patients^14-16^. The high prevalence of *RAS* mutations supports the notion that mechanisms of primary and acquired resistance are often similar. A small number of studies assessed acquired anti-EGFR-Ab resistance in tumor biopsies^11,17^. These also identified *RAS* and *EGFR* mutations but their retrospective nature and the analysis of only a small number of candidate genes may have biased the results. Increased growth factor availability, for example of ligands for the EGFR or MET RTKs^18,19^, confer anti-EGFR-Ab-resistance *in vitro* but their clinical relevance remains unknown. We undertook the first prospective trial that collected baseline (BL) biopsies immediately before initiation of single agent cetuximab and re-biopsies at the time of progressive disease (PD) from metastatic *RAS* wt CRCs. Biopsies were subjected to whole-exome mutation analysis, DNA-copy number analysis and to RNA-sequencing to assess primary and acquired resistance. Detailed insights into resistance mechanisms may enable more precise therapy allocation to patients who are likely to respond and open novel therapeutic opportunities for cetuximab-resistant CRCs.

## RESULTS

40 patients (pts) treated with single agent cetuximab could be assessed for treatment response and had sufficient biopsy material available for molecular analysis. Sequencing of BL biopsies failed in 5pts, leaving 35 for study (Patient characteristics and Consort-diagram: Supplementary fig.1 and table 1; Somatic mutation data: Supplementary table 2). As expected for CRC, this showed a high prevalence of *TP53* and *APC* mutations (Fig.1a). The median PFS and OS of this cohort were 2.6mo and 8.5mo, respectively (Supplementary fig.2a). 20pts showed primary progression at or before the first per-protocol CT scan (scheduled at week 12). The remaining 15 were classified as patients with prolonged clinical benefit. PD-biopsies were taken shortly after radiological progression (median: 14d after cetuximab cessation) from 25/35 cases and 24 were successfully exome sequenced. Sufficient RNA for RNA-sequencing was obtained from 25 BL- and 15 matched PD-samples.

**Figure 1:**
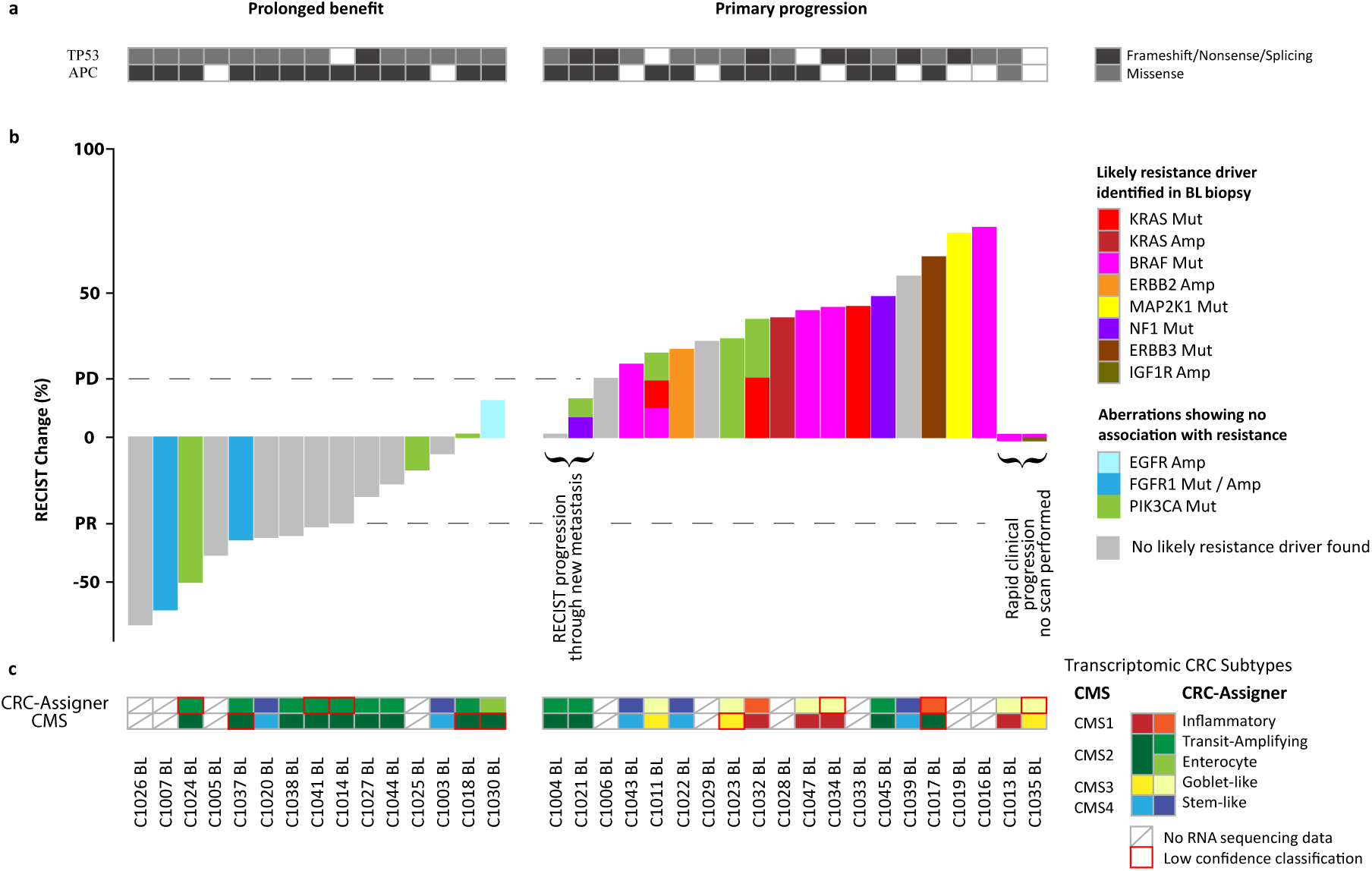
Molecular profiles of baseline (BL) biopsies categorized into cases with prolonged cetuximab benefit and primary progressors. **a:** TP53 and APC mutations detected in BL biopsies. **b:** Waterfall plot of best radiological response and genetic aberrations of RAS/RAF pathway members or regulators and PIK3CA detected by whole exome analysis. **c:** Transcriptomic subtype assignment. Amp=amplification, Mut=mutation. The figure legend for the transcriptomic subtypes is arranged to show the most similar CMS and CRCassigner subtypes next to each other and colored similarly.

### Genetic drivers of primary resistance

We first aimed to identify resistance drivers in BL-biopsies from 20 primary progressors (Fig.1b). Oncogenic *BRAF* V600E mutations were present in 6pts, one in combination with an *IGF1R* amplification (C1035BL, see Supplementary fig.3 for integer copy number profiles). No radiological response occurred in any of these and PFS was short (Supplementary fig.2b), supporting prior data that BRAF V600E confers resistance to single agent cetuximab^20^. C1011BL harbored a non-canonical *BRAF* D594F mutation, disrupting the DFG-motif of the kinase site. This is predicted to lead to a kinase-impaired BRAF variant^21^, which have been shown to paradoxically hyperactivate downstream ERK-phosphorylation when combined with oncogenic *RAS* alterations^22^. C1011BL indeed harbored a concomitant *KRAS* L19F mutation which has an attenuated phenotype compared to *KRAS* codon 12/13 hotspot mutations^23^. Another *KRAS* mutation (A18D) with an attenuated phenotype *in vitro* ^24^ was present on all 7 alleles of the polysomic chromosome 12p (Supplementary fig.4a) in C1033BL, likely explaining resistance in this case. A *KRAS* G12D mutation was identified in C1032BL, which had been found to be *KRAS* wt prior to study entry, indicating either a false negative result of the clinical assay or *KRAS* intratumor heterogeneity. A *KRAS* amplification was present in C1028BL and an *ERBB2* amplification in C1022BL (Supplementary fig.3). C1019BL harbored a canonical activating *MAP2K1/MEK1* mutation (K57N) and a concomitant *MAP2K1/MEK1* mutation (S228A) that did not influence kinase activity in a previous study^25^. Two tumors carried disrupting mutations in *NF1* (C1021BL: frameshift, C1045BL: nonsense). Both showed loss of heterozygosity of the *NF1*-locus on Chr17 (Supplementary fig.4b), constituting biallelic inactivation of this tumor suppressor gene. *NF1* encodes for neurofibromin, a negative regulator of KRAS, whose loss leads to EGFR-inhibitor resistance in lung cancer^26^. This suggests *NF1* inactivation as a novel driver of primary cetuximab resistance in CRC. The RTK *ERBB3* was mutated (P590L) in C1017BL but this amino acid change had no impact on *in vitro* growth in a previous study^27^, questioning whether it confers cetuximab resistance.

In contrast to prior studies^7,8^, neither *PIK3CA* nor *FGFR1* aberrations clearly associated with resistance: 4/20pts with primary progression harbored activating *PIK3CA* mutations (2xE545K, G364R, and H1047R concomitant with *PIK3CA* amplification (Supplementary fig.4c)) but also 3/15pts with prolonged benefit (2xV344G, H1047R). A tumor with a high level *FGFR1* amplification (C1037BL) and one with an *FGFR1* R209H mutation (C1007BL), previously reported in Kallmann syndrome^28^, had partial responses and prolonged benefit. An *EGFR* amplification was found in one tumor (C1030BL) and this associated with prolonged benefit as described^7^.

Together, oncogenic aberrations of *KRAS, BRAF, MAP2K1/MEK1, ERBB2* and *NF1* and *KRAS* mutations with attenuated phenotypes occurring concomitantly with polysomy of the mutant allele or with a kinase-impaired *BRAF* variant were identified in 14/20pts (70%) with primary resistance and one further tumor harbored an *ERBB3* mutation with unknown significance. No amplifications or oncogenic aberrations of RAS/RAF-pathway genes or RTKs that could explain resistance were identified in 5/20 (25%) primary progressors.

### Validation of transcriptomic CRC-subtypes as non-genetic predictors of cetuximab benefit

BL-biopsies for which RNA sequencing could be performed (n=25) were next assigned to recently defined transcriptomic CRC-subtypes using the CRCassigner-^10^ and the Consensus Molecular Subtype-classifications^29^ (Supplementary fig.5). There are strong similarities between subtypes of both classifications and 21/25 cases (84%) were assigned to matching subtypes, confirming robust performance (Fig.1c, see legend for subtype nomenclature and most similar subtypes). The Transit Amplifying (TA)-subtype has previously been associated with cetuximab sensitivity^10^ and was significantly enriched among cases with prolonged benefit (3.4-fold, p=0.017, Fisher’s exact test). The TA-subtype is most similar to the CMS2-subtype in the CRC Consensus Molecular Subtype-classification and the CMS2-subtype was equally enriched among patients with prolonged cetuximab benefit (2.9-fold enrichment, p=0.015 Fisher’s exact test). This validates the TA/CMS2-subtypes as non-genetic predictors of single-agent cetuximab benefit.

### Genetic drivers of acquired resistance

PD-biopsies from 14 metastases that radiologically progressed after prolonged clinical benefit were successfully exome sequenced (Fig.2a), including biopsies from two different progressing metastases in C1027. We first investigated genes with a known role in cetuximab resistance. Only one acquired *KRAS* mutation was found among these PD-biopsies (C1005PD: G12C). This clonally dominant mutation (Supplementary fig.3d) was accompanied by an *EGFR* mutation (G322S) which has not previously been described and whose relevance is uncertain in the context of a well characterized cetuximab resistance mutation in *KRAS*. One biopsy acquired a *KRAS* amplification (C1037PD). C1024PD acquired a clonally dominant *EGFR* mutation which has not previously been described (D278N), locating to the EGFR extracellular domain II^30^ but not affecting cetuximab binding epitopes. Expression of *EGFR* D278N in a CRC cell line did not confer cetuximab resistance and introduction into the 3T3 fibroblast line showed no evidence of constitutive EGFR phosphorylation (Supplementary fig.6), suggesting that this is a passenger mutation. No other *RAS*, *EGFR*, *BRAF* or *ERK* mutations or amplifications were detected in PD-biopsies.

**Figure 2:**
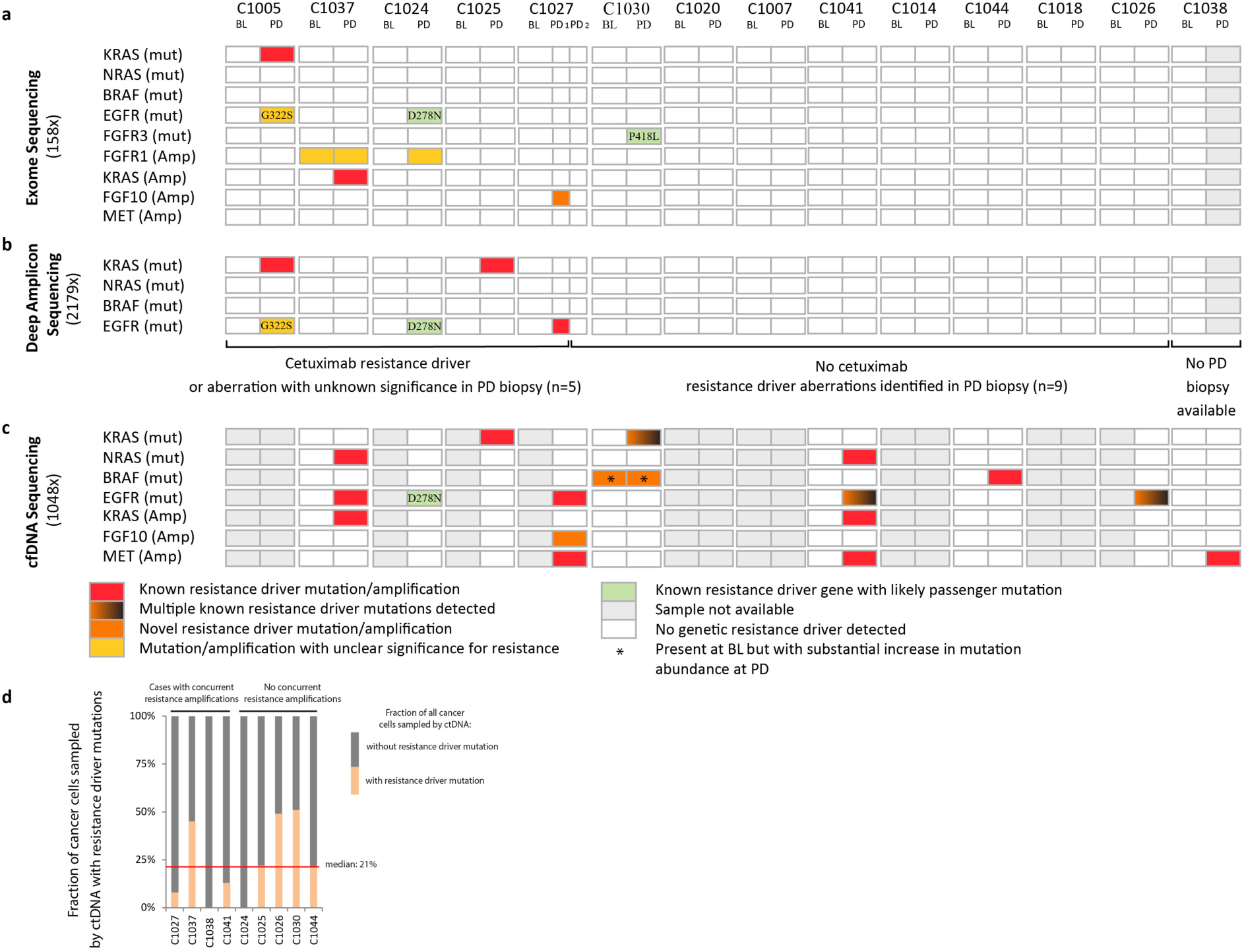
Genetic alterations acquired at progression (PD) in RAS/RAF pathway members or regulators. **a:** Mutations/amplifications identified by exome analysis of BL and PD biopsies. **b:** Ultra-deep amplicon re-sequencing of resistance driver hotspots in KRAS, NRAS, BRAF and EGFR in BL and PD biopsies. **c:** Mutations/amplifications identified by circulating tumor DNA sequencing **d:** Fraction of cancer cells sampled by ctDNA which harbored a resistance driver mutation at PD. The resistance fraction was calculated using VAFs of resistance driver mutations and the VAF of TP53 mutations, which is usually clonal/truncal in CRC, after correcting for the influence of local copy number status.

Two further RTKs acquired mutations at PD: *FGFR3* in C1030PD (P418L) and *ALK* in C1024PD (D626H). Neither located to the well defined mutational hotspots in these genes, nor have they been reported in the COSMIC cancer mutation database^31^, indicating that these may be passenger mutations. C1024PD furthermore acquired an *FGFR1* amplification (Supplementary fig.3). However, the presence of an *FGFR1* amplification in C1037BL which subsequently had a good response to cetuximab treatment (Fig.1b) questions whether this is sufficient to drive resistance. C1027PD1 acquired a narrow amplification (1.58Mbp, 60 DNA copies) encompassing *FGF10* (Supplementary fig.7a). *FGF10* encodes a ligand of the FGFR2 RTK which is expressed in most CRCs^32,33^. Recombinant FGF10 rescued growth and ERK-phosphorylation in CRC cell lines treated with cetuximab, supporting the notion that the acquired *FGF10* amplification is a novel driver of cetuximab resistance (Supplementary fig.7b). Different contributions of FGFR1 and FGFR2 to cetuximab resistance may result from differences in downstream signalling events^34^.

We next investigated genes that recurrently acquired mutations in PD-biopsies to investigate potential novel drivers of AR beyond the RAS/RAF-pathway. This revealed five large genes, each of which had acquired somatic mutations in two PD biopsies (Supplementary table 3). Based on reviews of gene functions and comparison of mutations against the COSMIC cancer mutation database^31^, none of these recurrently mutated genes were considered likely to confer cetuximab resistance (Supplementary discussion).

Acquired cetuximab resistance is often polyclonal^13^ and sequencing of PD-biopsies with a mean depth of 151x may have failed to detect resistance mutations in small subclones. We hence re-sequenced known cetuximab AR driver hotspots in *KRAS, NRAS, BRAF, MEK1* and *EGFR* by deep (2179x) amplicon sequencing in order to call potential subclonal mutations with VAFs as low as 0.5% (Fig.2b, Supplementary table 4). This revealed a *KRAS* Q61H mutation in C1025PD with a variant allele frequency (VAF) of 4.9%, and an *EGFR* exodomain S492R mutation with a VAF of 2.1% in C1027PD1. Both are known to confer cetuximab AR and were subclonal in these PD-samples (Supplementary fig.3).

Taken together, we identified known (*KRAS* mutations/amplifications, *EGFR* mutations of cetuximab binding epitopes) and novel (*FGF10* amplification) cetuximab resistance drivers in four PD-biopsies. One case acquired an *FGFR1* amplification and one an *FGFR3* mutation with unclear relevance for resistance. Importantly, no drivers of AR were found in 9/14 (64%) biopsied metastases despite each radiologically progressing (Supplementary Fig.8).

### Genetic drivers of acquired resistance in ctDNA

The low prevalence of cetuximab resistance drivers in PD biopsies was striking as it contrasts with results of ctDNA analyses of this trial and others that reported the evolution of *RAS* and *EGFR* aberrations in the vast majority of patients at the time they acquired cetuximab resistance^13,35^. We applied a ctDNA-sequencing assay targeting CRC and cetuximab resistance driver genes (Supplementary table 5) which simultaneously infers genome-wide copy number profiles^36^. This enabled us to correct VAFs for the influence of copy number states and to then quantify the proportion of the cancer cells that harbored resistance drivers by comparison against *TP53* mutations which are usually truncal in CRC^37^. Available ctDNA from 9 cases that progressed after prolonged cetuximab benefit (5 BL/PD pairs, 4 PD only) was deep sequenced (1048x mean depth). Known cetuximab resistance mutations in *RAS, BRAF* or *EGFR* were identified in 7/9 cases at PD (Fig.2c, Supplementary table 5). Furthermore, a kinase-impaired *BRAF* mutation (D594N) was detected in 6.8% of the cancer cell fraction in ctDNA at BL and this increased to 37.4% at PD in C1030 (Supplementary table 5). This clonal expansion during treatment and the identification of a kinase-impaired *BRAF* mutation in a primary resistant tumor (C1011BL) substantiates a role of these mutations in cetuximab resistance. DNA copy number profiles generated from ctDNA at PD furthermore identified amplifications of *MET* and *KRAS* in 3 and 2 cases, respectively (Fig.2c and Supplementary fig.9). The *FGF10* amplification found in the C1027PD1-biopsy was also identified in ctDNA at PD. Overall, ctDNA-sequencing revealed genetic drivers of AR in 8/9pts and frequent polyclonal resistance, similar to published ctDNA results^13^. We next used *TP53* mutations, detected in all ctDNA samples, to estimate the fraction of the cancer cell population represented in the ctDNA that harbored AR mutations at PD (Supplementary table 5). All detected resistance driver mutations taken together in each tumor were confined to a median of 21% of the cancer cells in the population (Fig.2d). The fraction of cancer cells that harbor an amplification cannot be estimated from ctDNA data as the absolute number of DNA copies in such subclones are unknown. Thus, only considering the five cases without concurrent AR amplifications in ctDNA, we still found a resistance gap with no detectable resistance mechanism in 49-100% of cancer cells sampled by ctDNA (Fig.2d). Although ctDNA and amplicon deep-sequencing may not identify very small subclones with genetic resistance drivers due to sensitivity limits, we hypothesized based on the ctDNA results and the inability to define genetic AR drivers in 9/14 biopsies (64%) from radiologically progressing metastases, that non-genetic resistance mechanisms may exist.

### Transcriptomic characteristics and their association with acquired resistance

The CMS2- and TA-subtypes significantly associated with cetuximab sensitivity at BL. Based on the observation that mechanisms of acquired drug resistance are often similar to those conferring primary resistance, we investigated whether subtype-identity may have a role in AR.

We first analysed PD-biopsies from tumors with prolonged benefit in which no genetic aberrations of cetuximab resistance genes had been found. Strikingly, 5/7 cases (71%) showed a switch from the cetuximab sensitive CMS2 subtype to the CMS4 subtype and 4/7 (57%) showed a TA to Stem-Like (SL) subtype switch (Fig.3a, Supplementary fig.5). In contrast, no CMS2/TA to CMS4/SL switches occurred in 6pts with primary PD. Switching from the cetuximab sensitive CMS2/TA-subtype to the CMS4/SL-subtype in the majority of PD-samples without identifiable genetic resistance mechanisms suggested that this contributes to acquired cetuximab resistance.

**Figure 3:**
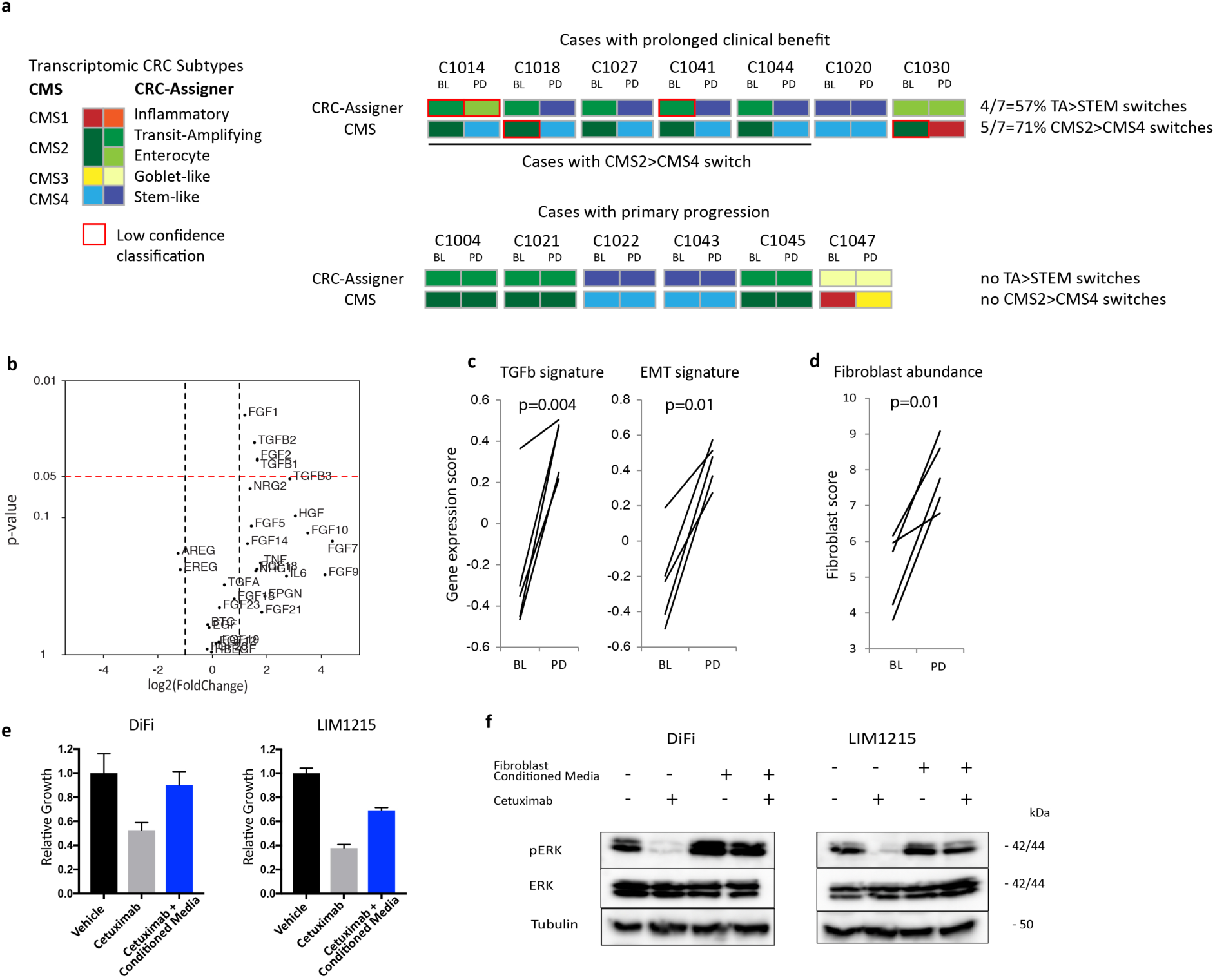
Transcriptomic CRC subtypes and fibroblasts as drivers of cetuximab resistance. **a:** Transcriptomic subtypes in baseline (BL) and progression (PD) biopsy pairs. **b:** Volcano plot showing differential expression of EGFR, MET and FGFR ligands in the five cases from **a** undergoing CMS2>CMS4 subtype switches at acquired resistance. **c:** Changes in TGFβ and EMT transcriptomic signatures in cases undergoing a CMS2>CMS4 subtype switch. **d:** Fibroblast abundance in cases undergoing a CMS2>CMS4 subtype switch based on MCPCounter analysis. **e:** Impact of CAF conditioned medium on the growth of CRC cell lines following 5-day cetuximab treatment with 50 µg/mL cetuximab. Error bars: standard deviation of six replicates. **f:** Western blot analysis showing CAF conditioned media rescue of pERK expression in CRC cell lines treated with 200 µg/mL cetuximab for 2 hours. p-values were calculated using the paired Student’s t-test.

Transforming growth factor beta (TGFβ)-expression is a defining characteristic of the CMS4/SL-subtypes. TGFβ1 and TGFβ2 RNA expression indeed significantly increased (3.1- and 2.9-fold increase in the means, p<0.05, paired t-test) in tumors undergoing a CMS2>4 switch (Fig.3b). TGFβ3 mean expression increased 7.2-fold at PD but this did not reach significance. A high level of TGFβ-activity in these samples was confirmed by the upregulation of a TGFβ-activated transcriptomic signature and of an epithelial to mesenchymal transition (EMT) signature which can be induced by TGFβ (Fig.3c).

CMS4 CRCs are enriched with cancer associated fibroblasts (CAFs) which are a major source of TGFβ and of mitogenic growth factors^38^. Applying the MCPCounter^39^ algorithm to RNA-sequencing data to bioinformatically assess the fibroblast content in the microenvironment confirmed significant increase in their abundance in PD-biopsies that had switched to the CMS4-subtype (Fig.3d). CMS2>4 subtype switches correspondingly increased the expression of growth factors that conferred cetuximab resistance *in vitro* (Fig.3b), including FGF1 and FGF2 (2.3- and 3.1-fold increase in the means, respectively) which activate multiple FGFRs and of the MET-ligand HGF which increased 8.3-fold, although the latter was not significant (p=0.1). In contrast, mean expression of the main EGFR-activating ligands in CRC, AREG and EREG, decreased 2.4-fold and 2.3-fold after subtype switching but this was not significant.

Conditioned media from cancer-associated fibroblasts (CAFs) can confer cetuximab resistance in colorectal cancer stem-like cells^40^. We questioned whether CAFs also promote resistance in well described cetuximab-sensitive CRC cell lines. Treatment with conditioned medium from immortalized CRC CAFs indeed rescued growth and maintained ERK phosphorylation in DiFi and LIM1215 CRC cell lines despite cetuximab treatment (Fig.3e and f).

Although this supports CMS2>CMS4 transitions and the associated increase in CAFs and mitogenic growth factors as a novel mechanism of acquired cetuximab resistance in CRC patients, two BL-biopsies from patients that subsequently achieved prolonged benefit from cetuximab therapy also displayed the CMS4-subtype. Thus, CMS4 identity does not invariably confer resistance. RNA-Sequencing data from BL- and PD-biopsies was available from one of these cases (C1020) and showed that TGFβ2 (4.4-fold), TGFβ3 (4.2-fold), HGF (2.7-fold) and FGF2 (1.6-fold) all increased from BL to PD (Supplementary table 6). This suggests a model where a gradual increase in growth factor-expression in a process associated with fibroblast infiltration and the acquisition of CMS4-subtype promotes resistance. This can evolve concurrently with genetic resistance in distinct subclones within the same patient, as demonstrated for cases that acquired CMS4 in a PD-biopsy while ctDNA showed the evolution of genetic resistance drivers in subclones (C1027, C1041, C1044). This parallel evolution of molecularly and functionally diverse resistance mechanisms within the same patients hinders the development of signalling pathway-targeting strategies to prevent or reverse resistance. The identification of new therapeutics that apply distinct selection pressures are hence a major need.

### Cetuximab impacts the cancer immune landscape

The IgG1-antibody cetuximab triggered immunogenic cell death and increased CRC immunogenicity in murine models^41^. Yet, whether effective cetuximab treatment promotes cancer immune responses in patients is unknown. We investigated this to explore potential opportunities to target cetuximab resistant CRCs with immunotherapy.

We first applied the cytolytic activity (CYT) signature^42^, which estimates the abundance of cytotoxic immune cells from RNA-expression data (Fig.4a). The mean CYT did not differ between BL-biopsies from tumors with prolonged benefit vs. those with primary progression (p=0.11, t-test). However, the mean CYT increased 5.9-fold from BL to PD in CRCs with prolonged benefit but not in those with primary progression, demonstrating that effective cetuximab treatment increased cytotoxic immune infiltrates. CYT remained relatively low in two tumors with prolonged benefit that showed no radiological shrinkage (C1018, C1030), suggesting that cancer cell death induction is required to stimulate cytotoxic cell infiltration. The largest CYT increases occurred in cases switching to the CMS4-subtype which is associated with an inflamed phenotype^29^. However, the median CYT in PD-biopsies of the five cases that switched to the CMS4-subtype was still 3-fold higher than in the five BL-biopsies classed as CMS4 prior to cetuximab exposure (Fig.4a). Hence, increased CYT after cetuximab therapy cannot be attributed to transcriptomic subtype changes alone.

**Figure 4:**
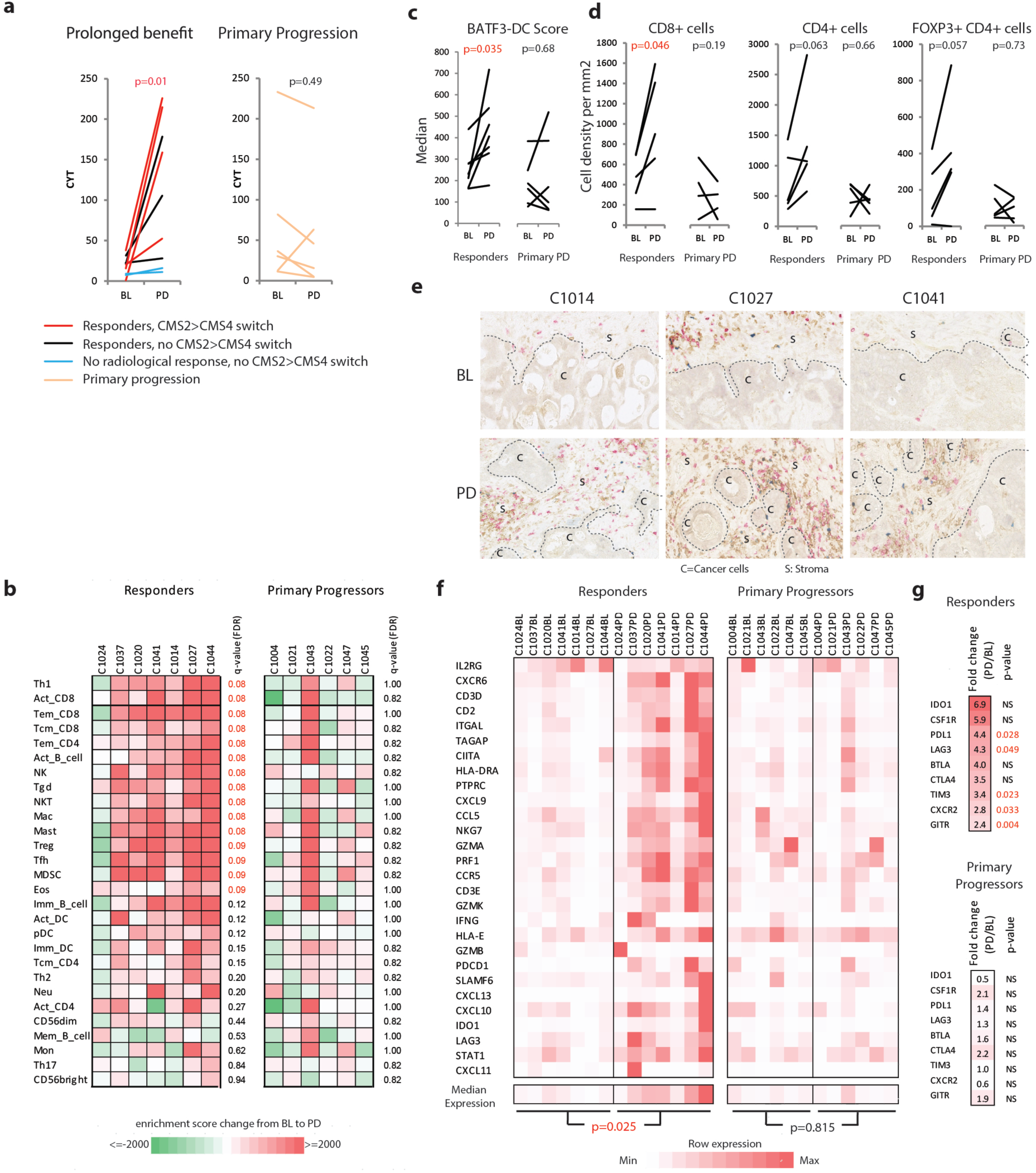
Impact of cetuximab on the tumor immune landscape. **a:** Cytolytic activity changes in paired BL and PD biopsies **b:** ssGSEA enrichment-score change from BL to PD samples for 28 immune cell subtypes. **c:** Transcriptomic score estimating the abundance of BATF3-positive dendritic cells **d:** Immuno-histochemical quantification of immune cell densities in FFPE specimens. **e:** Example of IHC of immune infiltrates before and after CMS2>CMS4 subtype switches (red: CD8, brown: CD4, blue: FOXP3, C=cancer cell area, S=stroma). **f:** Expression of a 28-gene signature of T-cell associated inflammation which has been predictive for immunotherapy outcomes across several cancer types **g:** RNA expression changes of targetable immune checkpoints and cytokine receptors. Statistical significance was assessed with the Mann-Whitney test followed by FDR correction (ssGSEA score changes) and the paired Student’s t-test.

Next, we bioinformatically inferred the abundance of 28 immune cell types from RNA-expression data^43^. A significant increase of T-cells that promote and execute adaptive immune responses, including all assessed CD8 T-cell subtypes, effector-memory CD4 and Th1 subtype T-cells, in PD-samples taken after cetuximab responses was observed (Fig.4b). Some immune cell subtypes which can dampen effective cytotoxic immune responses, including regulatory T-cells and myeloid-derived suppressor cells (MDSCs), also significantly increased. In contrast, immune cell infiltrates did not change in primary progressors. The presence of BATF3-positive dendritic cells (DCs) which cross-present antigens from dying cancer cells to CD8 T-cells is critical for immunotherapy efficacy in melanoma^44^. Applying a BATF3-DC score^45^ showed a significant increase at PD in tumors that had responded to cetuximab (1.7-fold increase, p=0.035) but no change in primary progressors (1.1-fold change, p=0.68, Fig.4c). Thus, several critical immune cell lineages for effective recognition of tumors by the adaptive immune system are enriched in tumors that responded to cetuximab.

To ascertain changes in immune infiltrates, we stained for CD8+, CD4+, and regulatory T-cells (FOXP3+ and CD4+) in paired BL and PD FFPE-biopsies available from 5pts with prolonged benefit and from 5 primary progressors (Fig.4d). CD8 T-cell densities increased significantly at PD compared to BL (2.0-fold change in means, p=0.047) in patients who responded to cetuximab. Mean of CD4+ and regulatory T-cells increased numerically but this was not significant (1.9-fold, p=0.057 and 2.2 fold, p=0.063), possibly because of the small number of cases available for this analysis. Thus, cetuximab treatment promotes T-cell infiltration of CRCs that respond and these are present at the time of progression. TGFβ has immunosuppressive effects, including the promotion of a T-cell excluded phenotype which can contribute to anti-PDL1 immunotherapy resistance^46^. Despite increased TGFβ expression after CMS2>CMS4 switching, CD8 T-cell infiltrates were present in intratumoral stroma in such PD-biopsies (Fig.4e), perhaps suggesting that the dynamic changes during response and re-growth prevent exclusion of T-cells.

We furthermore applied a signature of T-cell associated inflammation which is predictive for immune checkpoint-inhibitor benefit in several cancer types^47^. This significantly increased from BL to PD in responders but not in primary progressors (Fig.4f). Effective cetuximab therapy hence not only augments immune infiltrates including cytotoxic T-cells, but also T-cell associated inflammation which may indicate enhanced T-cell recognition of cancer cells. We finally questioned whether changes in immune infiltrates were accompanied by altered expression of immune-checkpoints or chemokine receptors which can be targeted by current immunotherapy agents. The immune-checkpoints LAG3, PDL1, TIM3 and GITR and the chemokine receptor CXCR2, which promotes myeloid cell infiltration, were significantly upregulated (Fig.4g). The up-regulation of immune-checkpoints may restrain T-cell infiltrates and could provide opportunities to develop novel therapeutic strategies following cetuximab failure.

## DISCUSSION

This prospective trial revealed novel associations of biallelic *NF1* loss and of non-canonical *RAS/RAF*-aberrations with primary resistance to single agent cetuximab. While *KRAS* A18D and L19F, and *BRAF* mutations other than V600E were rare in large CRC cohorts (each <1%)^48,49^, *NF1* mutations have been reported in ∼5% of cases and successful validation as a predictive marker in randomized trials could spare these patients ineffective treatment. Our results are supported by a study describing an association of *NF1* mutations with poor PFS with cetuximab in combination with chemotherapy^50^, although 3 out of 4 were missense mutations with unknown effects on *NF1* function and there was no testing for loss of heterozygosity.

In contrast to previous reports^7,51^, neither *PIK3CA* mutations nor *FGFR1* aberrations clearly associated with primary resistance. Particularly *PIK3CA* exon 20 mutations have been described to confer resistance to anti-EGFR-Ab in combination with chemotherapy, however we found the exon 20 mutation H1047R in a responder but also in combination with a *PIK3CA* amplification in a primary progressor. Concomitant copy number aberrations or the use of single agent cetuximab may explain these differences. The small sample size furthermore warrants cautious interpretation of these results.

We found a strikingly lower frequency of acquired resistance driver mutations in *RAS* and *EGFR* in PD-biopsies than anticipated based on the pervasive detection of these drivers in cetuximab treated patients analyzed by ctDNA^52^. The absence of genetic aberrations of cetuximab resistance driver genes in 64% of PD-biopsies, corroborated by ctDNA analysis which did not detect resistance drivers in 45-100% of the cancer cell population sampled, highlights a significant unexplained resistance gap. The majority of PD biopsies without identifiable genetic resistance drivers no longer displayed the cetuximab-sensitive CMS2/TA-subtype found before treatment initiation but the CMS4/SL-subtype which is rich in fibroblast and in growth factors that confer cetuximab resistance *in vitro*. Together with the inability to identify genetic resistance drivers in these biopsies from radiologically progressing metastases strongly suggests that subtype switching and associated stromal remodelling is a novel mechanism of acquired resistance to single agent cetuximab. This could explain similar genetic results in a series of 37 PD biopsies which found no aberrations in *RAS, BRAF or EGFR* in 46% of biopsies with acquired anti-EGFR-Ab resistance^53^.

These data demonstrate the limitations of ctDNA analysis which is restricted to the identification of genetic resistance mechanisms and the importance of parallel tissue analyses with multi-omics approaches. They furthermore portray a cetuximab resistance landscape resembling that of EGFR-inhibitors in lung cancer or BRAF-inhibitors in melanoma where non-genetic resistance can occur. Lung cancers that acquired resistance but showed no genetic resistance drivers can upregulate growth factors which activate bypass signalling pathways or EMT^54-56^ as alternative resistance mechanisms and fibroblast-mediated stromal remodelling can confer acquired BRAF-inhibitor resistance in melanomas^57^.

Combinatorial drug treatment approaches to reverse acquired cetuximab resistance in CRC appear challenging, due to toxicities that will most likely arise when attempting to combine multiple signalling-pathway inhibitors and because of the inability to effectively target *RAS*-mutant clones which evolved in 4/9pts. However, strategies to delay resistance by preventing subtype switching, for example by targeting TGFβ, a master-regulator of the CMS4/SL-subtype, could be assessed.

Our analysis of the immune landscape in CRCs that responded to cetuximab and then progressed shows significantly increased cytotoxic T-cells but also of immune-suppressive cells, such as regulatory T-cells and MDSC. This was accompanied by the upregulation of an inflammation signature which has been predictive of checkpoint-inhibitor success in other cancer types, potentially indicating a role for immunotherapy. The significant upregulation of immune-suppressive checkpoints such as PDL1 and LAG3 highlights testable strategies. Exploring immunotherapies after cetuximab resistance has been acquired may circumvent the limited clinical opportunities to directly target the frequently polyclonal and heterogeneous cetuximab resistance mechanisms.

## Online methods

### Trial design and samples

The Prospect-C trial (ClinicalTrials.gov identifier: NCT02994888), is a prospective translational study investigating biomarkers of response or resistance to anti-EGFR-Ab-therapy in *KRAS* wt chemo-refractory metastatic CRC. No *NRAS* mutant cases were enrolled as the licensed cetuximab indication changed to *KRAS and NRAS* wt CRC during the trial. Patients who were at least 18 years old and had a World Health Organization performance status of 0-2, were eligible if: all conventional treatment options including fluorouracil, irinotecan, oxaliplatin were exhausted or patients were intolerant/had contraindications for oxaliplatin/irinotecan-based chemotherapy; they had metastatic cancer amenable to biopsy and repeat measurements with computed tomography (CT) scanning.

Written informed consent was obtained from all patients. The study was carried out in accordance with the Declaration of Helsinki and approved by the national UK ethics committee (UK Research Ethics Committee approval: 12/LO/0914). All participants were required to have mandatory image-guided pre-treatment biopsies (targeted to the CT identified index lesion), and mandatory biopsies at the time of RECIST-defined progression (from one or two suitable progressing metastatic sites). Treatment consisted of single-agent cetuximab at a dose of 500mg/m^2^ administered every other week until progression or intolerable side effects.

The identification of novel biomarkers of primary and acquired resistance to cetuximab therapy in DNA and RNA from CRC tumor biopsies was the primary endpoint of the study. The study recruited to the recruitment target of 30 patients that had been treated and had BL and PD samples available for genetic analyses. After removing cases with insufficient DNA yield or tumor content based on sequencing results, data from 24 paired BL and PD samples was available for mutation and copy number analysis. 11 cases from which only a BL biopsy was available were included in the analysis. Secondary endpoints included the identification and validation of novel biomarkers for resistance and response to cetuximab in RNA and ctDNA. The trial protocol also permitted further exploratory molecular analyses.

The efficacy parameters including partial response and stable disease were measured using RECIST v1.1 criteria. Progression free survival (PFS) was measured from start of treatment to date of progression or death from any cause. Overall survival (OS) was defined as time from start of treatment to death of any cause. Patients without an event were censored at last follow up before PFS and OS were estimated.

The cohort was dichotomized into primary progressors who had PD before or on the first per protocol CT scan, scheduled at 12 weeks from the start of cetuximab treatment. This was performed at a median of 12 weeks with a range of 9-16 weeks on treatment. Patients with prolonged benefit were defined as those who remained progression free at the time of this scan.

Samples from healthy donors were collected for ctDNA sequencing after obtaining written informed consent through the ‘Improving Outcomes in Cancer’ biobanking protocol at the Barts Cancer Centre (PI: Powles), which was approved by the UK national ethics committee (Research Ethics Committee approval: 13/EM/0327).

### Sample preparation

DNA and RNA were extracted simultaneously from snap frozen biopsies using the Qiagen All Prep DNA/RNA Micro Kit following the manufacturer’s instructions. Matched normal DNA was extracted from blood samples using the Qiagen DNA Blood Mini Kit. DNA concentration was measured using the Qubit dsDNA Broad Range Assay Kit, and integrity checked by agarose gel electrophoresis. A minimum quantity of 500 ng, and where available 2 µg of DNA, was used for next generation sequencing. RNA from biopsies which were successfully DNA sequenced was subjected to RNA-Sequencing if a sufficient quantity (>125ng) and quality (RIN>5.5) was confirmed by electrophoresis on the Agilent 2100 Bioanalyzer. Blood for circulating tumor DNA analysis was collected in EDTA tubes and centrifuged within 2 hours (10min, 1600g) to separate plasma, which was stored at −80°C. Upon thawing, samples were further centrifuged (10min, 16000g, 4°C). ctDNA was extracted from up to 4 ml plasma per patient and from 2×4 ml from healthy donors using the Qiagen QIAamp Circulating Nucleic Acid Kit. ctDNA was quantified on the Agilent 2100 Bioanalyzer.

### Whole exome/genome DNA sequencing

Biopsy samples were sequenced by the NGS-Sequencing facility of the Tumour Profiling Unit at the Institute of Cancer Research (ICR) or at the Beijing Genome Institute (BGI). Exome sequencing libraries were prepared from a minimum of 500 ng DNA using the Agilent SureSelectXT Human All Exon v5 kit according to the manufacturer’s protocol. Paired-end sequencing was performed on the Illumina HiSeq 2500 platform with a target depth of 100X for exomes (ICR) and on the Illumina HiSeq X10 platform with 70X for genomes (BGI).

### Bioinformatics analysis of DNA sequencing data

BWA-MEM^58^ (v0.7.12) was used to align the paired-end reads to the hg19 human reference genome to generate BAM format files. Picard Tools (http://picard.sourceforge.net) (v2.1.0) *MarkDuplicates* was run with duplicates removed. BAM files were coordinate sorted and indexed with SAMtools^59^ (v0.1.19). BAM files were quality controlled using GATK^60^ (v3.5-0) *DepthOfCoverage*, Picard *CollectAlignmentSummaryMetrics* (v2.1.0) and fastqc (https://www.bioinformatics.babraham.ac.uk/projects/fastqc/) (v0.11.4).

### Somatic mutation analysis

Tumor and germline DNA sequencing results were assessed for matching SNP profiles to check for potential sample swaps. This identified one case where germline DNA and tumor DNA SNP profiles differed and this was removed from the analysis. For single nucleotide variant (SNV) calls we used both MuTect^61^ (v1.1.7) and VarScan2^62^ (v2.4.1). SAMtools (v1.3) *mpileup* was run with minimum mapping quality 1 and minimum base quality 20. The pileup file was inputted to VarScan2 *somatic* and run with a minimum variant frequency of 5%. The VarScan2 call loci were converted to BED file format and BAM-readcount (https://github.com/genome/bam-readcount) (v0.7.4) run on these positions with minimum mapping quality 1. The BAM-readcount output allowed the VarScan2 calls to be further filtered using the recommended *fpfilter.pl* accessory script^63^ run on default settings. MuTect was run on default settings and post-filtered for minimum variant allele frequency 5%. Indel calls were generated using Platypus^64^ (v.0.8.1) *callVariants* run on default settings. Calls were filtered based on the following FILTER flags - ‘GOF, ‘badReads, ‘hp10,’ MQ’, ‘strandBias’,’ QualDepth’,’ REFCALL’. We then filtered for somatic indels with normal genotype to be homozygous, minimum depth >= 10 in the normal, minimum depth >=20 in the tumor and >= 5 variant reads in the tumor. Exonic regions were analyzed in whole genome sequenced samples to assure comparability to the whole exome sequenced samples. Mutation calls were further filtered with a cross-normal filter by running bam-readcount on the bed file of merged variants for all sequenced matched normal (blood) samples. For both SNV and Indel calls we used a threshold of >=2% of the total number of reads at the call loci. If the alternate allele count is equal to or greater than this threshold the variant is flagged as present in the normal sample. A call is rejected if the variant is flagged in 5% or more of the normal samples in our cohort to remove common alignment artifacts or those arising recurrently at genomic positions which are difficult to sequence.

Mutation calls were merged and annotated using annovar^65^ (v1.8.0.0) with hg19 build version. The allele counts were recalculated using bam-readcount with minimum base quality 5 (in line with minimum default settings of the joint SNV callers). The calls were then filtered on minimum variant allele frequency >=5%, minimum depth >=20 in a called sample and a maximum of 2 variant alleles in the matched normal sample.

### DNA copy number aberration analysis

CNVKit^66^ (v0.8.1) was run in non-batch mode for copy number evaluation. We first identified high confidence SNP locations using bcftools^59^ (v1.3) *call* with snp137 reference and snpeff ^67^ (v4.2) *SnpSift* to filter heterozygous loci with minimum depth 50. We further extracted positions spaced 500bp apart in the whole genome samples. VarScan2 was used to call the tumor sample BAMs at these locations to generate B-Allele Frequency (BAF) data as input for CNVKit.

We generated basic *access* and *antitarget* files to indicate the accessible sequence regions. This excluded blacklisted regions suggested by CNVKit and the HLA region. We then generated a pooled normal sample and used the *winsorize* and *pcf* functions within copynumber^68^ to identify further outlier positions and regions of highly uneven coverage. These regions were merged to ensure consistency across all data.

CNVKit was run with matched normals along with the adjusted access and antitarget files. For the segmentation step we ran *pcf* from the R-package copynumber. Breakpoints from this segmentation step were then fed into Sequenza^69^ (v2.1.2) to calculate estimates of purity/ploidy and these values were used as a guide to recenter and scale the LogR profiles in CNVKit. BAF and LogR profiles were also manually reviewed by two researchers to determine their likely integer copy number states. Adjustments were made in cases where both manual reviews identified a consensus solution that differed from the bioinformatically generated integer copy number profile. Furthermore, BL/PD sample pairs where the ploidy of one sample was close to double the ploidy of the other sample and copy number profiles were highly similar (suggestive of a genome doubling event), the sample with lower ploidy was adjusted to a the likely genome-doubled higher state to facilitate a direct comparison of copy number changes, unless clear evidence of BAF and LogR profiles suggested otherwise. These adjustments were made in samples C1004PD, C1022PD, C1025PD, C1027PD1, C1030PD, C1043BL where both manual reviews supported a different solution to Sequenza.

### Analysis of gene amplifications

Amplifications were defined as a 3-fold or greater increase on the ploidy of a sample, a substantial loss event as a 3-fold or greater decrease on the ploidy state and a homozygous deletion as CN=0. Amplification and loss threshold values were rounded to the nearest integer copy number state. Ploidy was estimated as follows,

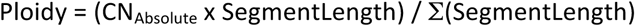

with CN_Absolute_ representing the unrounded copy number estimate and SegmentLength the genomic length between segment break points. BL and PD biopsy pairs were compared to identify which cases had acquired amplifications at PD that were absent at BL.

### Deep amplicon sequencing

Ampliseq libraries were prepared by the ICR-TPU using the Ion Chef from 800 ng DNA extracted from BL/PD biopsies, and from matched germline samples. A custom amplicon panel comprising a single pool of 77 amplicons (Supplementary Table 4 for amplicon positions) was designed to cover mutational hotspots and known cetuximab resistance drivers in KRAS, NRAS, BRAF, EGFR and MAP2K1 and several mutations identified by exome sequencing in each sample (including any TP53 and APC mutations) to enable subclonality estimates. Up to 32 samples were pooled and sequenced on PGM 318 chips (v2) with 500 flows. Deep amplicon sequencing data was aligned and somatic mutations were called using the Ion Torrent Suite software (v5.2.2). run with a minimum variant frequency of 0.5% and 3 supporting variant reads.

### ctDNA-sequencing

Ultra-deep circulating tumor DNA (ctDNA) sequencing with molecular barcode error correction^36^ was applied to cases with prolonged benefit from cetuximab and which had at least 25 ng of ctDNA. Libraries were prepared from 25 ng ctDNA using the Agilent SureSelect^XT-HS^ kit and hybridized to a CRC panel targeting up to 40 genes (Supplementary table 5) using our optimized protocol^36^. Libraries were pooled and sequenced on an Illumina HiSeq2500 in 75bp paired-end mode, generating a median of 125.7M reads/sample.

The resulting data was aligned and molecular barcode-deduplicated in order to reduce false positive sequencing errors using Agilent SureCall, with variants called using the integrated SNPPET caller. To call very low frequency variants, bam-readcount was used to interrogate targeted hotspot positions in *KRAS, NRAS, BRAF, ERK1 (MAP2K1)* and *EGFR* (detailed in Supplementary table 3). In order to maximize the sensitivity for the detection of resistance mutations, these were called if at least 2 independent variant reads were identified at a mutational hotspot position and encoded for a recurrently observed amino acid change in the specific gene. Genome-wide copy number profiles were constructed using CNVKit run in batch mode with Antitarget average size 30 kb as described^36^. ctDNA sequenced from healthy donors^36^ was used as the normal reference dataset. Copy number profiles generated from ctDNA were aligned with copy number profiles showing absolute copy numbers from matched biopsies and the closest integer copy number was assigned to TP53 and mutated cetuximab resistance driver genes for the subclonality analysis.

### RNA-sequencing

NEB polyA kit was used to select the mRNA. Strand specific libraries were generated from the mRNA using the NEB ultra directional kit. Illumina paired-end libraries were sequenced on an Illumina HiSeq2500 using v4 chemistry acquiring 2 × 100 bp reads. Bcl2fastq software (v1.8.4, Illumina) was used for converting the raw basecalls to fastqs and to further demultiplex the sequencing data.

Tophat2 spliced alignment software^70^ (v2.0.7) was used to align reads to the GRCh37 (hg19) release-87 human reference genome in combination with Bowtie2^71^ (v2.1.0). FeatureCounts^72^ was used to perform read summarization. Sample QC was performed using Picard Tools *CollectRnaSeqMetrics*. We excluded 2 samples (C1006BL and C1007BL) with fewer than 10% of reads aligning to exonic regions. Lowly expressed genes were filtered using a cpm threshold equivalent to 10/L, where L is the minimum library size in millions^73^. Sample batch effects were assessed using principal component analysis and did not require corrective action. Counts were normalized for library size using *estimateSizeFactors* in Deseq2^74^. FPKM data were generated using the *fpkm* function in Deseq2. For downstream analysis all data were filtered for protein coding genes using the GTF annotation file and filtering on the *gene_biotype* column.

### Statistical analyses

All statistical analysis was performed using R (v3.4.0) and STATA13. The Fisher’s exact test was used to examine association of categorical variables in 2×2 contingency tables. The Student’s ttest was applied to examine means of continuous data (e.g. normalized RNA-Sequencing counts, cytolytic activity scores, median expression values of the T cell associated inflammation signature, immunohistochemical immune cell densities and MCP-counter^39^ fibroblast infiltrate scores from non-paired sample groups). The paired Student’s t-test was applied to these datasets when comparing paired (BL and PD) data. p-values ≤0.05 were considered statistically significant. The Kaplan-Meier method was used to estimate OS and PFS probability. The Mann-Whitney statistical test was applied to compare ssGSEA rank scores of 28 immune cell populations followed by False Discovery Rate correction and a q value ≤ 0.1 was considered statistically significant.

### Cancer cell content analysis

The cancer cell content of each sequenced sample was assessed based on the variant allele frequency (VAF) of somatic mutations and samples with an estimated cancer cell content below 10% were removed from the analysis as the sequencing depth was insufficient to accurately detect mutations in these samples^61^. As the majority of mutations are heterozygous and hence present in half of the DNA copies of the cancer cells, 2xVAF can be used to approximation the fraction of cancer cells in a sample. This led to the exclusion of four samples (C1001BL, C1009BL, C1010BL, C1042BL) as shown in the Consort diagram (Supplementary fig.1a). The median estimated cancer cell content across the remaining 60 samples was 41% (Supplementary table 1).

### Subclonality analysis exome sequencing data

The clonal status of mutations was assessed using the allele specific copy number generated in the CNVKit solution. We estimated the cancer cell fraction (CCF) using the phyloCCF method as described by Jamal-Hanjani et al.^75^. We then inferred the mutation copy number (i.e. the number of alleles harbouring the mutation) and assigned clonal/subclonal status to each variant using the criteria described by McGranahan *et al.*^76^.

### Subclonality analysis in ctDNA and amplicon sequencing data

Variant allele frequencies of *TP53* mutations, of hotspot resistance driver mutations in *KRAS, NRAS, BRAF* and *EGFR* and of the *EGFR* mutation D278N were extracted from ctDNA BAM files. *TP53* mutation VAFs were used to calculate what fraction of the ctDNA was of cancer cell origin by correcting for the influence of copy number aberrations using the following formula:

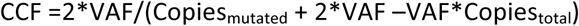

with CCF indicating the cancer cell fraction, Copies_mutated_ the number of copies that harbored the *TP53* mutation and Copies_total_ the absolute copy number of the *TP53* locus. Clonality analysis of TP53 mutation showed clonal mutations and loss of heterozygosity of the TP53 locus for all tumour biopsies with the exception of C1027 which harbored two TP53 mutations, one present on four copies of chromosome 17p and one on two copies, suggesting biallelic inactivation through two distinct mutation events. TP53 Copies_mutated_ and Copies_total_ were equal for tumors with TP53 LOH and in 1027 the VAFs of both TP53 mutations were taken together and the sum of all chromosome 17p copies were used to estimate CCF.

The same formula was then resolved to calculate the expected VAF of a clonal mutation given the CCF of the ctDNA sample and the local copy number state of this mutation:

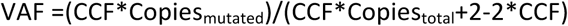

Copies_total_ for all mutations were inferred from ctDNA copy number profiles that had been close matched to the integer copy number states of biopsies (Supplementary fig 9). For subclonality calculation, we furthermore assumed that resistance drivers were only mutated on a single gene copy (i.e. Copies_mutated_=1, which is likely as they are thought to have a dominant effect). This assumption furthermore maximized the estimated fraction of cancer cells that harbor a resistance driver mutation, hence providing a conservative measure of the resistance gap. The fraction of the total CCF in ctDNA that harbors an observed resistance driver mutation was then calculated by dividing the observed VAF by the expected VAF for a mutation that is 100% clonal. We then estimated the maximum fraction of all cancer cells that harbored resistance driver mutations in a sample as the sum of the CCF values of all individual resistance driver mutations in that sample.

### Colorectal cancer subtyping

Consensus Molecular Subtypes^29^ were assigned using CMScaller^77^. The CMScaller function was run with raw count data and setting ‘RNASeq=TRUE’. Each sample was assigned the subtype with the shortest distance according to the inbuilt nearest template prediction (NTP)^78^. The CMScaller classification was considered low confidence if FDR >0.01. Samples were also assigned to the molecular CRC subtypes, as described by Sadanandam et al.^10^. To minimize technical differences in subtype assignment we generated data normalized using the same approach as CMScaller (limma::normalizeQuantiles(log2(x+.25))). The data were then row median centered and correlated with the PAM centroids, as defined by the published 786-gene CRCAssigner signature. Each sample was then assigned to the CRC subtype with the highest correlation. If the correlation coefficient is <0.15 or the difference with the second highest coefficient is <0.06 then the sample is considered low confidence^29^. The EMT and TGFβ expression signatures were generated by the Camera Gene Set Analysis in CMScaller for each sample.

The subtyping showed a high level of agreement between the classification approaches. This was true even of assignments considered low confidence by the published criteria.

### Immune cell infiltrate analysis

The cytolytic activity (CYT) was calculated as the geometric mean of the *GZMA* and *PRF1* genes (normalized expression values as input, offset by 1.0). The BATF3-DC signature was calculated as the mean of the normalized expression values of the genes in this signature. FPKM normalized RNA sequencing data and published immune cell metagenes^43^ were used as input into the single sample gene set enrichment analysis (ssGSEA) algorithm using default settings to determine immune cell enrichments in each sample as described^79^.

The Microenvironment Cell Populations (MCP)-counter algorithm^39^ was used as an independent bioinformatics tool to assess immune cell enrichment. Data were normalized using limma^80^ *voom* and the MCPCounter function run with *HUGO_symbols* chosen as *featuresType*.

### Immunohistochemistry

5 µm slides were cut from FFPE blocks and triple stained as described^81^. 5 representative tumor areas of 0.05 mm^2^ were identified per slide and CD8+ cells, FOXP3+ and CD4+ cells and CD4+ FOXP3- were quantified in each of the selected areas at x40 magnification using ImageJ software. Densities were calculated as cells/mm^2^. Immune cell scoring was performed blinded.

### Cell lines

DiFi and LIM1215 cell lines were a gift from the Valeri Lab at ICR. NIH-3T3 cells were a gift from the Huang Lab at ICR. DiFi cells were cultured in RPMI-1640 (Gibco), GlutaMax (Gibco), 5% FBS. LIM1215 cells were cultured in RPMI-1640, 10% FBS, hydrocortisone (Gibco), 1-thioglycerol (Sigma) and insulin (Gibco). NIH-3T3 cells were cultured in DMEM (Gibco), GlutaMax (Gibco) and 10% FBS. Human fibroblasts from rectal carcinomas which have been immortalized using hTERT virus (pCSII vector backbone) (RC11) were a gift from Fernando Calvo, initially provided by Danijela Vignjevic (Institute Curie, France)^82^. Fibroblasts were cultured in DMEM (Sigma), GlutaMax (Gibco), 10% FBS, 1% insulin-selenium-transferrin (ITS, Gibco). Conditioned media was harvested from flasks of confluent RC11 cells after 72h.

### Drug Assays

FGF10 rescue experiments were performed in DiFi and LIM1215 colorectal cancer cell lines treated with cetuximab and FGF10 (Peprotech) for 7 days. Treatments were replenished with fresh media after 3 days. EGFR mutant experiments were performed in LIM1215 cells. Cells were treated with cetuximab for 5 days. DiFi and LIM1215 cells were seeded in standard media or CAF-conditioned media and treated with cetuximab for 5 days. All experiments were perfumed in six replicates. Viability was assessed using CellTiter Blue reagent (Promega) for all assays.

### DNA constructs

The Gateway Entry clone R777-E053-Hs.EGFR was a gift from Dominic Esposito (Addgene plasmid #70337). Site directed mutagenesis using QuikChange Lightning (Agilent) and custom designed primers (5’-CCTCGGGGTTCACATTCATCTGGTACGTGGT/5’-ACCACGTACCAGATGAATGTGAACCCCGAGG) was used to generate the EGFR-D278N mutant. The fulllength sequence was assessed using Sanger sequencing to confirm presence of the intended mutation and that no other mutations had been inserted before Gateway recombination into the lentiviral expression construct pLX304 (a gift from David Root, Addgene plasmid #25890). HEK293T cells were infected with the lentiviral constructs pLX304-WT and pLX304-D278N in combination with packaging plasmids psPAX and pMD2.G (a gift from Didier Trono, Addgene #12260 and #12259 respectively). LIM1215 and NIH-3T3 cells were transduced with the resultant viral supernatants in the presence of Polybrene (8 µg/mL), and selected with 0.5 µg/mL Blasticidin.

### Western Blotting

Total cell lysates were prepared using NP-40 buffer supplemented with protease and phosphatase inhibitors (Sigma). Samples were resolved by electrophoresis on SDS-PAGE gels for Western blotting. Primary antibodies used were p-ERK (Cell Signalling Technologies #9101), ERK (Cell Signalling Technologies #9102), p-EGFR (Cell Signalling Technologies #2236) and EGFR (Cell Signalling Technologies #2232). Bands were detected using HRP-labelled secondary antibodies and ECL Prime, GE Healthcare), followed by visualisation on an Azure Biosystems C300 detection system.

## Acknowledgements

The study was supported by the National Institute for Health Research Biomedical Research Centre for Cancer at the ICR/RMH and by the CRUK Immunotherapy Incubator Grant to the ICR, RMH and UCL. MG, AW, LB and BG were supported by CRUK, a charitable donation from Tim Morgan, Cancer Genetics UK and the Constance Travis Trust. GS was funded by an Institute of Cancer Research Studentship. RGE was supported by a grant from the Spanish Society of Medical Oncology (FSEOM) for Translational Research in Reference Centers. The ICR Centre for Evolution and Cancer was supported by a Wellcome Trust Strategic Grant (105104/Z/14/Z). We thank all patients, relatives and the clinical trials unit and tissue collection team who made this study possible.

## Author contributions

KhK coordinated the Prospect C trial. AW and MND performed bioinformatics analysis and MG, MND, FS, KK, LJB and GS analyzed the data. BG and GS performed *in vitro* analyses, LJB performed ctDNA and amplicon sequencing. SAQ, TM and AF performed and RGE and KvL analysed immunohistochemistry stains. MG conceived, funded and supervised the molecular analysis. DC is the chief investigator of the Prospect C trial and obtained funding for the trial. SG assessed protein changes for functional significance. RR and FC immortalized, characterized and provided the CAF line. AS and YP supported the molecular subtype analysis. NS, IC, SR and DW recruited treated patients in the Prospect C trial which was supported by the clinical research team (RB, IR, JT, AB, AW, NK, NF). CP, KyK and AW performed statistical analysis. The pathologists KvL and AW analyzed and interpreted tissue sections. MG, AW, LJB, GS and KhK wrote the manuscript and all authors approved the final manuscript.

